# mtDNA recombination indicative of hybridization suggests a role of the mitogenome in the adaptation of reef-building corals to extreme environments

**DOI:** 10.1101/462069

**Authors:** Eulalia Banguera-Hinestroza, Yvonne Sawall, Abdulmohsin Al-Sofyani, Patrick Mardulyn, Javier Fuertes-Aguilar, Heiber Cárdenas-Henao, Heiber Cárdenas-Henao, Francy Jimenez-Infante, Christian R. Voolstra, Jean-François Flot

## Abstract

mtDNA recombination following hybridization is rarely found in animals and was never until now reported in reef-building corals. Here we report unexpected topological incongruence among mitochondrial markers as evidence of mitochondrial introgression in the phylogenetic history of *Stylophora* species distributed along broad geographic ranges. Our analyses include specimens from the Indo-Pacific, the Indian Ocean and the full latitudinal (2000 km) and environmental gradient (21°C-33°C) of the Red Sea (N=827). The analysis of *Stylophora* lineages in the framework of the mitogenome phylogenies of the family Pocilloporidae, coupled with analyses of recombination, shows the first evidence of asymmetric patterns of introgressive hybridization associated to mitochondrial recombination in this genus. Hybridization likely occurred between an ancestral lineage restricted to the Red Sea/Gulf of Aden basins and migrants from the Indo-Pacific/Indian Ocean that reached the Gulf of Aden. The resulting hybrid lives in sympatry with the descendants of the parental Red Sea lineage, from which it inherited most of its mtDNA (except a highly variable recombinant region that includes the *nd6*, *atp6*, and mt*ORF* genes) and expanded its range into the hottest region of the Arabian Gulf, where it is scarcely found. Noticeably, across the Red Sea both lineages exhibit striking differences in terms of phylogeographic patterns, clades-morphospecies association, and zooxanthellae composition. Our data suggest that the early colonization of the Red Sea by the ancestral lineage, which involved overcoming multiple habitat changes and extreme temperatures, resulted in changes in mitochondrial proteins, which led to its successful adaptation to the novel environmental conditions.

## 1. INTRODUCTION

Studies of mitochondrial genomes have been of broad relevance for understanding the ecological and evolutionary processes leading to the diversification of organisms (Ballard and Rand 2005; Chinnery and Hudson 2013; Kivisild 2015; Wolff et al. 2016), due particularly to key characteristics such as: maternal inheritance (in most organisms); high mutation rates in triploblastic metazoans (but not in cnidarians and poriferans; cf. Shearer et al. 2002); and lack of recombination (Barr et al. 2005). However, the latter assumption has been challenged by the increasing evidence of mtDNA recombination in animals (Rokas et al. 2003; Barr et al. 2005; Tsaousis et al. 2005; Ujvari et al. 2007; Ladoukakis and Zouros 2017) with important ecological and evolutionary implications (Rokas et al. 2003; White et al. 2008; Dokianakis and Ladoukakis 2014; Levsen et al. 2016).

In natural populations, most evidence for mtDNA and/or nuclear introgression comes from species living at the edge of their geographic range (i.e. marginal ecosystem that often exhibit genetically atypical populations; Johannesson and André, 2006), perturbed habitats, near extreme environmental conditions or in hybrid zones (Riginos and Cunningham 2004; Johannesson and Carl 2006; Brennan et al. 2014; Taylor et al. 2015; Hellberg et al. 2016). There is growing evidence in plants, in fungi, and to a lesser extent in animals, that divergent lineages that meet and interbreed in contact zones, particularly after expanding their range as a consequence of climatic or habitat fluctuations, produce hybrids carrying recombinant mitochondrial genomes (Barr et al. 2005) that are able to adapt to new environmental or habitat conditions leading eventually to hybrid speciation (Riginos and Cunningham 2004; Brennan et al. 2014; Taylor et al. 2015; Mastrantonio et al. 2016).

Marginal areas such as the Red Sea, a biogeographic *cul-de-sac* extending from the Indian Ocean, are observed to host a high number of hybrid species (Veron 1995; Johannesson and Carl 2006; DiBattista et al. 2016; Berumen et al. 2017) and therefore offer an interesting playground to study the main factors influencing patterns of diversification and speciation in extreme environments. This basin exhibits a high variability in oceanographic conditions following a N-S gradient along four well-defined oceanographic provinces (Raitsos et al. 2013), ranging from relatively low temperatures in the northern areas (below 23°C) to record summer temperatures up to 33°C in the southern regions (Acker et al. 2008; Raitsos et al. 2013; Moustafa et al. 2014; Sawall et al. 2014; Sawall et al. 2015).

In addition, since its inception this region has undergone a complex climatic and geological history. During the last glacial cycles, the Red Sea experienced multiple sea-level fluctuations (Siddall et al. 2003; Rohling et al. 2013) as well as repetitive periods of connection and isolation from the Indo-Pacific, through the Strait of Bab-el-Mandeb in the South (Righton et al. 1996; Siddall et al. 2004; Rohling et al. 2013). These events, together with discontinuous periods of extinction and re-colonization from refuge zones inside and outside the region (Fine et al. 2013; Pellissier et al. 2014), likely played an important role in shaping the geographic distribution of genetic diversity for coral reefs in this oceanic basin, as well as the diversification of their associated fauna (DiBattista et al. 2013; DiBattista et al. 2016a; DiBattista et al. 2016b; Berumen et al. 2017). In fact, it has been postulated that corals entered the region during the Miocene –a period characterized by hyper-saline conditions and strong climate change– and after several episodes of post-Miocene migrations (Taviani 1998). Therefore, lineages already inhabiting this area were affected by the successive geological and climatic processes influencing the region (DiBattista et al. 2016). Repetitive invasions from the Indian Ocean likely enhanced the potential for the encounter of lineages with different genetic backgrounds, increasing the chance for hybridization and therefore hybrid speciation in the unique conditions of this marine zone.

Introgressive hybridization has been reported to be widespread in reef-building corals, with records mostly in the families Acroporidae, Poritidae and Pocilloporidae (Van Oppen et al. 2001; Willis et al. 2006; Combosch and Vollmer 2015; Richards and Hobbs 2015; Hellberg et al. 2016; Forsman et al. 2017; Johnston et al. 2017). Indeed, the topological incongruence observed in coral phylogenies might be explained by reticulate evolution due to multiple climatic and see-level changes across geological eras (Veron 1995; Van Oppen et al. 2001; Vollmer and Palumbi 2002; Willis et al. 2006). Moreover, it has been proposed that many species occurring in marginal areas and with restricted distribution, such as those endemic of the Indo-Pacific Ocean, might be hybrids (Veron 1995; Richards and Hobbs 2015). Even though mtDNA introgression is expected to be more frequent than nuclear introgression, particularly between species occurring in sympatry (Mastrantonio et al. 2016), it has not being reported to date in reef-building corals.

The coral genus *Stylophora* (family Pocilloporidae) is widely distributed in the Indo-Pacific and Indian Ocean and comprises a range of a highly plastic morphospecies (Veron 1995; Veron 2002), which are diversified within four phylogenetic lineages (Flot et al. 2011; Stefani et al. 2011; Keshavmurthy et al. 2013). *Stylophora* is found throughout the entire environmental range of the Red Sea and is represented by several morphospecies: *Stylophora pistillata* (Esper, 1797), *Stylophora subseriata* (Ehrenberg, 1834), *Stylophora wellsi* (also found in Madagascar), *Stylophora kuelhmanni*, *Stylophora danae* and *Stylophora mamillata* (Scheer and Pillai 1983; Veron 2002). A recent molecular study aiming to identify species boundaries in the genus (Arrigoni et al. 2016) proposes that all these morphospecies belong to one single cohesive molecular lineage.

We selected the genus *Stylophora* given its widespread geographical distribution and the presence of several endemic morphospecies in the Red Sea (Veron et al. 2018), which may represent important sources of genetic variability for the future adaptation of reef-building corals under the current threats of climate changes. Coral species within this genus and their associated zooxanthellae were studied across the environmental gradients of the Red Sea, spanning over ~ 2000Km (12 degrees of latitude) and sequences from other relevant biogeographic regions were also included (i.e. New Caledonia, Great Barrier Reef, Madagascar, Gulf of Aden and Arabian Gulf). A total of 827 sequences were analyzed to understand their genetic diversity, phylogenetic history and phylogeographic framework, in addition 281 colonies were typified for *Symbiodinium*.

Our study was centered in mtDNA genes, but nuclear markers were included to understand differences in phylogenetic and phylogeographic patterns. Molecular analyses included ITS2 for *Symbiodinium* and four nuclear (*hsp70*, ITS1, ITS2 and *PMCA*) and five mitochondrial genes for the coral host (mtCR, *cox1*, 12S, 16S and *atp6*-mt*ORF*). Subsequently, the patterns of diversification of the Red Sea specimens, outlined by mtDNA genes, were evaluated using full mitochondrial genome sequences of species belonging to the family Pocilloporidae (*Pocillopora*, *Seriatopora*, *Stylophora* and *Madracis*) (Veron and Pichon 1976).

Our aims were (1) to investigate patterns of diversification in *Stylophora* species along broad geographic ranges and verified further whether host morphological variation had any taxonomic value for Red Sea species (in other words, is there any association between morphological and genetic variation?). (2) to find out whether mitochondrial variation, zooxanthellae association, and phylogeographic patterns fit the trends expected from the environmental gradients in this marginal basin, and (3) to test whether the topological incongruence between mtDNA markers observed in our data and previously reported by Arrigoni et al. (2016) could be explained by introgressive hybridization. Here we discuss our results from an integrative perspective, coupling phylogenetic and phylogeographic patterns, *Symbiodinium* distribution, morphospecies-clades association and the finding of mtDNA recombination. We also discuss the implications of these results for coral conservation in highly variable environments.

## 2. RESULTS

### 2.1. Gene variation and polymorphic sites

Our largest data set consisted of 827 sequences of the mt*ORF* locus, which includes a short region of the adjacent *atp6* gene, from a large portion of the geographic distribution of *Stylophora* (Figure 1, Supplementary Table *S1*); it included 716 samples from our collection of the Red Sea, 3 samples from the Arabian Gulf, plus 108 sequences of *S. pistillata* morphospecies from Stefani et al. (2011) and Flot et al. (2011): from the Gulf of Aden (N=20), Pacific regions (N=60) and Madagascar (N=28). In addition, we produced a second dataset for four other mitochondrial fragments, from CR (N=401), *cox1* (N=147), 12S (N=72) and 16S (N=61) and four nuclear fragments, from *hsp70* (N=738), ITS1 (N=75), ITS2 (N=56) and *PMCA* (N=86) genes. This data set includes sequences from other studies (see material and methods section 4).

**Figure 1.**
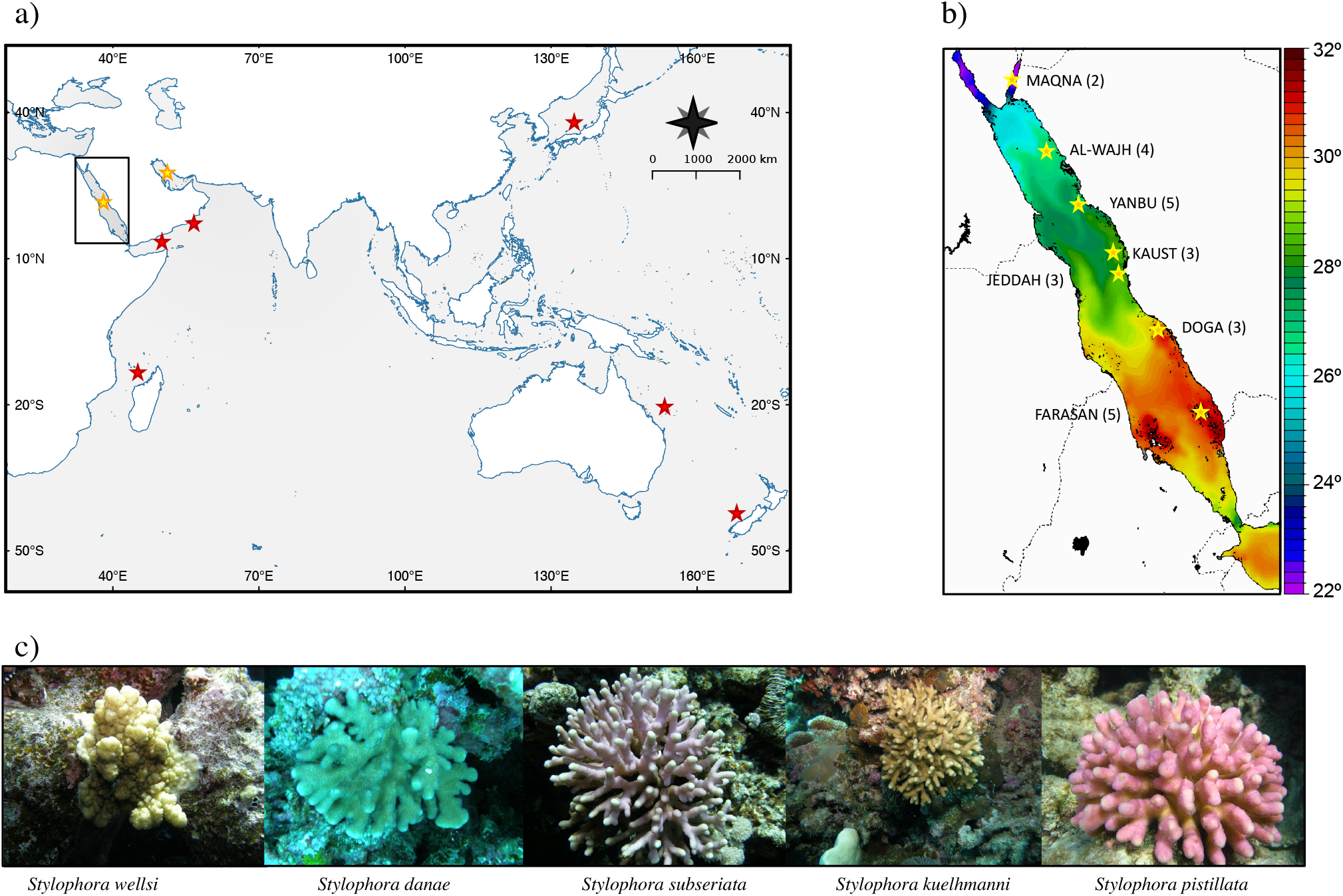
Sampling localities and *Stylophora* morphotypes. **a)**. General map showing the sampling sites. Red stars indicate sequences or samples obtained from published work and yellow stars specify samples collected for this study. **b)**. Map indicating sampling sites along the environmental gradient of the Red Sea. The numbers in parenthesis show the reefs visited at each location. The map was modified from the original downloaded from https://www7320.nrlssc.navy.mil/global_ncom/glb8_3b/html/Links/red/temp_glb8_3b_2012060700_0000m.gif. **c)**. *Stylophora* morphotypes included in this study.

While the *atp6-*mt*ORF* locus (N=827; 884-943 bp) displayed a high level of polymorphism in *Stylophora* samples, the two *cox1* fragments (303-545 bp) were found monomorphic when comparing sequences across all Red Sea specimens. Pairwise comparisons among all individuals uncovered multiple synonymous and non-synonymous substitutions at the 5’ end and 3’ end of the mt*ORF* gene, but noticeably, the core region was composed mainly by duplicated tandem repeats, which were of variable lengths and dissimilar among *Stylophora* specimens from the Red Sea (in terms of nucleotide composition and length of the repeat; data not shown).

In addition, the analysis of the full *atp6* sequences revealed a high number of mutations in this gene. Two haplotypes were found among Red Sea specimens differing by 50 point mutations, which were differentiated from an Indo-Pacific/Indian Ocean haplotype in 45 and 8 point mutations respectively. Most changes at the *atp6* gene were synonymous, except two changes found at positions 104 (i.e. Asparagine –AAT– / Valine –GTT– / Isoleucine –ATT–) and at position 187 (Serine –TCA–/Phenylalanine –TTT–/ Leucine –TTA–) resulting in changes that involved amino acid with different polarities.

The CR (N=401; 757 bp) was the most variable of the non-coding markers, with polymorphism mainly caused by variation in the number of tandem repeats, some of them being specific to *Stylophora* specimens inhabiting different geographic areas. Polymorphic sites, indels and parsimony informative sites for mitochondrial and nuclear genes are recorded in Supplementary Table *S2*.

### 2.2. Phylogenetic analyses

ML and Bayesian phylogenies inferred from the *atp6*-mt*ORF* locus including *Pocillopora* as an outgroup (no shown) or excluding the outgroup (Figure 2a) showed a large phylogenetic subdivision between two groups, each associated with both high Bayesian Posterior Probabilities (BPP > 1.0) and high bootstrap values (> 99%). A first clade, (N=461; h=27) included some of the sequences from the Red Sea and the Gulf of Aden and those from the Arabian Gulf (N=373; h=12), plus sequences from Madagascar (N=28, h=6) and the Pacific (N=60; h=9); the second clade included the other specimens from the Red Sea and Gulf of Aden (N=366; h=14) –we refer to this clade as the Red Sea Lineage B (*RS_LinB*)– (Figure 2a). These two lineages were also evident in the mitogenome phylogenies of the family using the full *atp6* (678 bp) and *nd6* genes (564 bp) (Figure 3a) and in the phylogenies derived from our recombination analyses (Figure 3b) –See recombination results below–.

**Figure 2.**
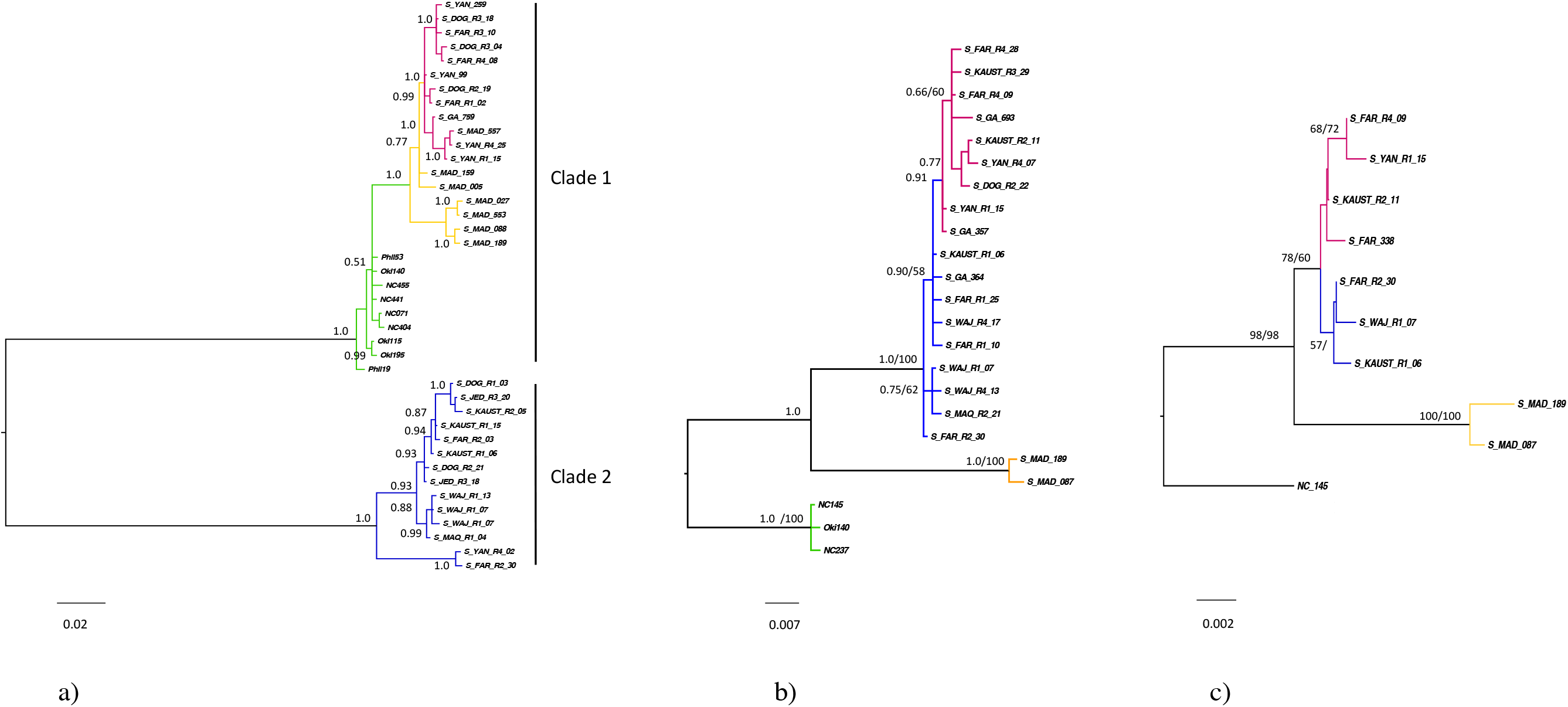
Mitochondrial phylogenies of the genus *Stylophora*. **a)**. Bayesian tree of the *ATP6*-mt*ORF* locus. **b)**. Bayesian tree for the CR gene. **c)**. Maximum Likelihood (ML) tree for the 12S gene. Node support are given in Bayesian Probabilities (BPP) for the mt*ORF*, BPP/ML for the mtCR and ML/NJ for the 12S. Colors are as follow: Red: Haplotypes from Arabian Gulf, Red Sea and Gulf of Aden placed in Clade 1 (*RS_LinA*). Yellow: haplotypes from Madagascar. Green: haplotypes from Indo-Pacific (Philippines, Okinawa, New Caledonia) and Blue: Red Sea and Gulf of Aden haplotypes placed in Clade 2 (*RS_LinB).* Haplotypes names are given in agreement to the region where the haplotype was found in higher frequency (MAQ=MAQNA, WAJ=ALWAJH, YAN=YANBU, KAUST = KAUST, JEDDAH=JED, DOG = DOGA, FARASAN = FAR)

**Figure 3.**
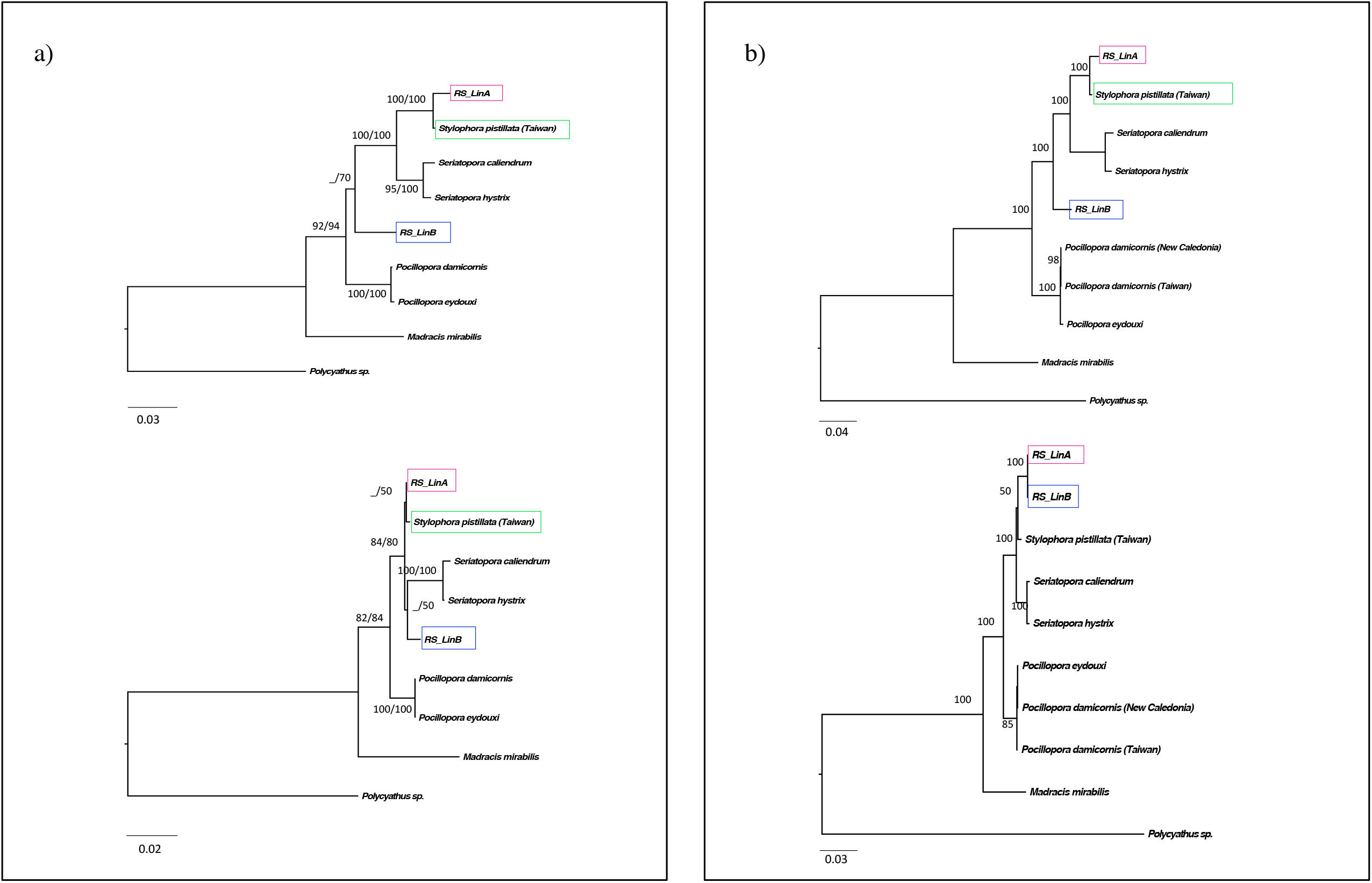
Mitochondrial phylogenies of the family Pocilloporidae and phylogenies derived from the recombination analyses. **a)**. Position of the *RS_LinA* and *RS_LinB* in a family phylogeny of the full *atp6* (above) and *nd6* genes (below). **b)**. ML phylogenies from the recombination hypotheses: phylogeny derived from the recombinant region –minor parent–, including the *nd6*, *atp6*, and *mtORF* genes; 2415bp (above) and phylogeny from mtDNA genes outside the recombinant region –from the major parent –, 9555bp (below).

In contrast, in the phylogenetic trees inferred from all other markers, whether mitochondrial (CR, 12S) or nuclear (ITS1; ITS2*, hsp70; PMCA*) (Figures 2b and Supplementary Figure *S1*), *RS_LinA* and *RS_LinB* sequences form a clade that exclude all sequences from Madagascar and from the Indo-Pacific region. There is therefore a major incongruence between the phylogenetic signal of *atp6*, *nd6* and the mt*ORF* vs the CR and 12S (Figure 2 and Figure 3a) and between mtDNA genes vs the nuclear genes. The ITS2 showed a pattern similar to that of the CR, in which *RS_LinA* and *RS_LinB* are almost completely separated; there is only one individual from *RS_LinB* placed inside the clade formed by *RS_LinA* sequences, and vice versa, on the ITS2 tree (Supplementary Figure *S1*). The *cox1* and the 16S trees provided only little phylogenetic information (not shown).

### 2.3. Comparative analyses between *RS_LinA* and *RS_LinB*

#### 2.3.1. Intraspecific phylogenies and morphospecies-clade association

Bayesian and ML trees obtained based on the full-length sequences of the *atp6*-mt*ORF* locus, thus analyzing members of each lineage (*RS_LinA*, 884 bp, h=33; *RS_LinB*, 943 bp, h=20) separately, are shown in Figure 4. In general, we found no clear association between Red Sea morphospecies and phylogenetic groups in the *RS_LinA* lineage. For example, some mitochondrial haplotypes were shared among different morphotypes or/and individuals exhibiting the same morphotypes were placed into different subclades (Figure 4a); the only exceptions were haplotypes belonging to morphotypes of *S. kuelhmanni*, that were all associated with subclade *LA1*, and those from *S. mamillata –*from Arrigoni et al. (2016)– that were exclusively found in subclade *LA3* (data not shown). Instead, in the *RS_LinB* lineage, morphotypes attributed to *S. subseriata* (called *Stylophora* cf. *subseriata* thereafter) were grouped in subclade *LB1* and displayed haplotypes found exclusively in the northern Red Sea – see phylogeographic analyses below– and were differentiated from morphotypes belonging to *S. pistillata* (mainly short and medium branched morphotypes, called as *Stylophora* cf. *pistillata* from now on), whose haplotypes were distributed mainly in the central-southern Red Sea and were placed in subclade *LB2* (Figure 4b).

**Figure 4.**
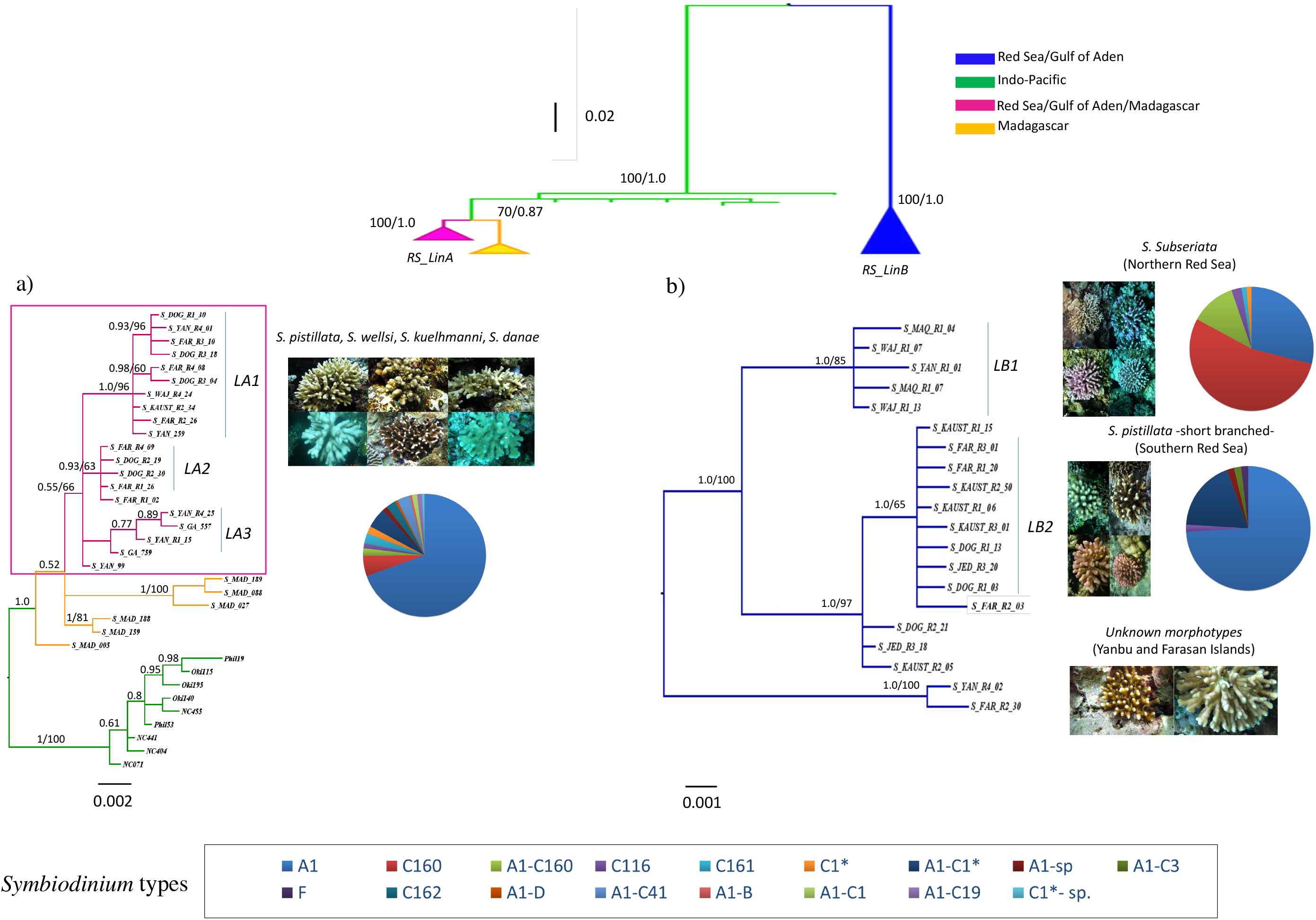
Extended *atp6*-mtORF phylogenies for **a)**. Clade 1. Highlighting the *RS_LinA* lineage (magenta) and the different subclades within this group. **b)**. Clade 2. Exclusively composed by Red Sea/Gulf of Aden specimens (*RS_LinB*). The two subclades within *RS_LinB* correspond with northern (colder) and central-southern (hotter) regions. The relationship among haplotypes, morphotypes, and *Symbiodinium* types are shown for each lineage.

#### 2.3.2. *Symbiodinium* distribution

As for zooxanthellae, *Symbiodinium* type A1 (*Symbiodinium microadriaticum*) was the most abundant in members of *Stylophora* within the Red Sea (167 out of 281 colonies), followed by *Symbiodinium* type C160 (N=49). The distribution of these types was somehow different between members of each *Stylophora* mtDNA lineage (Figure 4). In colonies placed within the *RS_LinB* and inhabiting the northern Red Sea (i.e. subclade *LB1*; *Stylophora* cf. *subseriata*; N=76) the most abundant type was C160 (N=41) followed by type A1 (N=22) and few colonies presented both types (A1-C160; N=9). In contrast, colonies distributed in the central-southern areas (i.e. subclade *LB2*; i.e. *Stylophora* cf. *pistillata*) were mainly associated with type A1 (40 out of 54 colonies) and the presence of C160 type, or of this type in combination with A1 (A1-C160) were not detected. In southern regions, few colonies exhibited the C1* type, which was not present in northern colonies and was always found in association of type A1 (i.e. A1-C1*; N=10). Other C types or associations A1-C types were found in very low abundance (i.e. in one colony). A high percentage of the colonies of the *Stylophora RS_LinA* (105 colonies out of 151) showed association with *Symbiodinium microadriaticum* in both northern and southern regions. The few colonies carrying C160 or A1-C160 types were mainly found in Yanbu (7 out of 11 colonies). The presence of C1* and A1-C1* types were detected mostly in colonies inhabiting the southernmost area of the Red Sea (i.e. Farasan; 7 out of 8 colonies). Few C types (i.e. C161, C162, C116) and combination of A1 with other types (i.e. A1-D, A1-C41, A1-B, A1-C1, A1-C19; A1-sp) displayed low frequencies and some of them were exclusive of colonies belonging to this group (Figure 4).

### 2.4. Comparative phylogeography of *RS_LinA* and *RS_LinB* and patterns of genetic variation along the Red Sea gradient

Median-joining networks were constructed for the markers for which we had the largest number of sequences: *atp6*-mt*ORF* (N=827), CR (N=401) and *hsp70* (N=738). Each network indicates a clear phylogeographic signal (i.e., there was a visible association between genetic variation and geographic location). The north-south genetic differentiation was more pronounced for *RS_LinB* than for *RS_LinA*, a pattern supported by all three loci (Figure 5). Haplotype frequencies along the gradient were somehow different among lineages. In *RS_LinB* the highest frequencies of the most common haplotypes were found either in northern or southern areas, while haplotype frequencies in *RS_LinA* showed a clinal variation (Supplementary Figure *S2*). For the *hsp70*, haplotypes showed different frequencies along the gradient when evaluated per mtDNA lineage (Supplementary Figure *S2c*) and the pattern was similar to that shown by the *atp6*-mt*ORF* locus.

**Figure 5.**
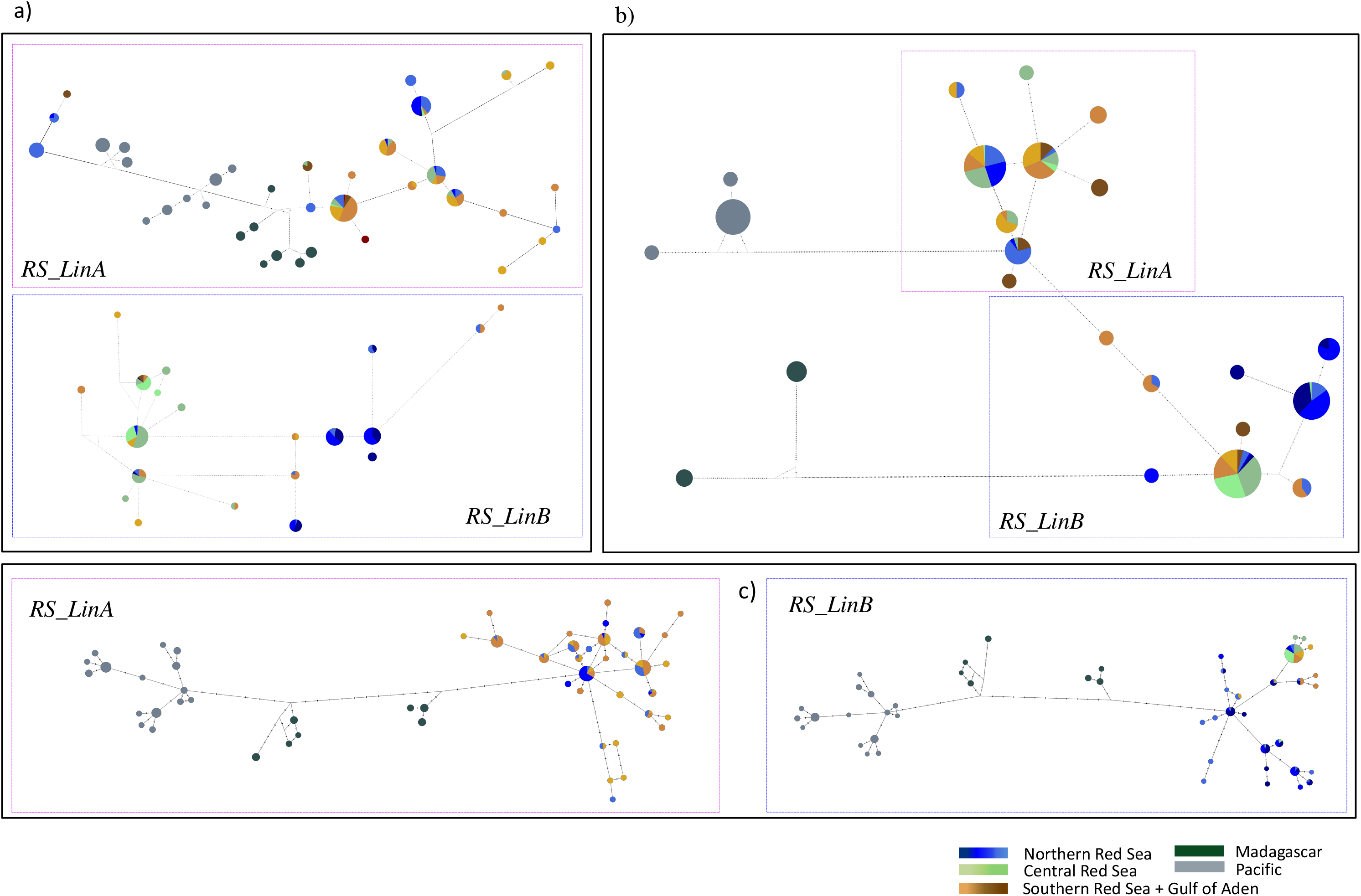
Network trees showing the relations among haplotypes for the three more variable genes. **a)**. *atp6*-mt*ORF*. **b)**. Control Region (CR) **c)**. *hsp70*. Vertical lines indicate the number of mutations.

### 2.5. mtDNA recombination analyses

The analyses of the mitogenomes for all members of the family Pocilloporidae including *RS_LinA* and *RS_LinB* and the genus *Polycyathus*, resulted in the detection of several recombination events in pocilloporid corals. Recombination signals were detected in *Stylophora, Seriatopora and Madracis.* The signals for *Stylophora* and *Seriatopora* were confirmed when excluding the two divergent genomes of *Madraci*s and *Polycyathus*. In *Stylophora* the strongest signal supported by all seven methods was found in *RS_LinA*, with the major parent hypothesized as the *RS_LinB* and the minor parent *Stylophora pistillata* (Taiwan). *P*-values were as follow: RDP = 1.175 × 10^−10^; GENECONV = 8.96 × 10^−11^; BOOTSCAN = 1.237 × 10^−11^; MAXCHI = 7 × 10^−16^; CHIMAERA = 4.019 × 10^−03^; SISCAN = 1.090 × 10^18^ and 3SEQ = 2.969 × 10^−29^. The boundaries of the breakpoints were placed at the end of the *nd2* genes by all methods, extending across the *nd6*, *atp6*, mt*ORF* and ending at the beginning of the *nd4* gene (Supplementary Figure *S3*). A second event, in which the recombinant region was transferred from *Pocillopora* to the *RS_LinB* was hypothesized, in this case the recombinant region was restricted to the *atp6* gene. A third event was detected, by only 3 methods, in *Seriatopora hystrix and Seriatopora caliendrum*.

Family phylogenies focused in *Stylophora* and including the region inherited from the minor parent (*S. pistillata*, Taiwan; 2415 bp) and the major parent (*RS_LinB*; 9555 bp) (Figure 3b) corroborated the phylogenetic patterns outlined for *RS_LinA* and *RS_LinB* in our interspecific phylogenies (Figure 2).

## 3. DISCUSSION

The results from our study show the first evidence of mitochondrial recombination in corals and support the hypothesis that mtDNA introgressive hybridization is the most likely cause of topological mito-mito incongruence in *Stylophora*. Our recombination analyses, using the full mitogenomes of pocilloporid corals (*Madracis, Pocillopora, Seriatopora* and *Stylophora*, including the two Red Sea lineages), provides evidence of mtDNA introgression in this group – particularly in *Stylophora–* strongly suggesting the existence of a hybrid lineage (*RS_LinA*) in the Red Sea, Gulf of Aden and Arabian Gulf.

The mitogenome of *RS_LinA* contains introgressed genes from two divergent mitochondrial lineages: on mtDNA genes found in the recombinant region (i.e. covering the complete *nd6*, *atp6* and mt*ORF* genes), which have been inherited from a parental species included in a lineage distributed in the Indo-Pacific/Indian Ocean region, while all other mtDNA genes can be traced back to the ancestor of the *RS_LinB*. In the Red Sea, the *RS_LinA* is found in sympatric association with the descendants of its local parental species (*RS_LinB)*, and they harbour different phylogeographic patterns and different *Symbiodinium* composition. In addition, we found that morphospecies placed within the *RS_LinB* agreed with phylogenetic subclades, which accounts for the presence of two divergent allopatric populations divided in northern and southern areas, consistent with environmental and oceanographic discontinuities, a trend that was completely absent in the hybrid *RS_LinA*.

The colonization of the extreme environments of the Red Sea by the ancestral *RS_LinB* implied the development of adaptations to demanding temperature and salinity conditions. This likely involved selective pressures observed to multiple amino acid changes in mitochondrial proteins, evidenced by the highest substitution rate in the *atp6* and in the mt*ORF* genes of the *RS_LinB*, when compared with other *Stylophora* lineages. For example, the *atp6* gene showed a high number of nucleotide substitutions in the ancestral *RS_LinB*, when compared with the hybrid and the Indo-Pacific/Indian Ocean lineage (i.e. 50 and 45 point mutations respectively), and only few substitutions between the hybrid and its parental Indo-Pacific species (i.e. 8 mutations), which suggest a role of such mtDNA genes in the adaptation to extreme environments. These results have important implications for the conservation of Red Sea species and corals.

### 3.1. Divergent mitochondrial lineages of *Stylophora* are present in the Red Sea

Our *atp6*-mt*ORF* phylogenetic analyses highlight the presence of two divergent sympatric *Stylophora* lineages in the Red Sea/Gulf of Aden/Arabian Gulf areas: the *RS_LinA* sharing a common ancestry with an Indo-Pacific/Indian Ocean/Madagascar lineage –placed in Clade 1– and the *RS_LinB* lineage, which form a unique clade on its own –Clade 2– (Figure 2). The existence of these two lineages was reinforced by the family phylogenies of the full *atp6* and *nd6* genes (Figure 3a), which support the hypothesis that the hypervariable mt*ORF* gene, a mitochondrial open reading frame of unknown function, which is unique to the family Pocilloporidae (Flot and Tillier, 2007a), perform as a barcode gene for members of this family (Johnston et al. 2017).

Noticeable, the *atp6*-mt*ORF* showed topological incongruence with the CR/12S loci (Figures 2b, 2c) and the four nuclear genes used in these analyses did not support the existence of *RS_LinA* and *RS_LinB* lineages (Supplementary Figure *S1*). The only nuclear gene that showed a similar phylogenetic pattern of that shown by the CR and 12S genes, was the ITS2, which likely imply that the ITS2, subject to concerted evolution, tells a similar evolutionary history to that of mitochondrial genes.

Mito-nuclear incongruence such as reported in this study have already been reported in the literature on many metazoans taxa (Willis et al. 2006; Toews, Brelsford 2012; Pavlova et al. 2013; Forsman et al. 2017; Salvi et al. 2017) and they are frequently invoked as one of the main signatures of selection (Morales et al. 2015), the result of incomplete lineage sorting (Van Oppen et al. 2001; DeBiasse et al. 2014; Richards and Hobbs 2015) or introgressive hybridization (Diekmann et al. 2001; Gompert et al. 2008; Forsman et al. 2017). The latter is often considered to be rampant in corals, with reticulate evolution having been proposed as one of the main processes that contributed, in great extent, to their successful adaptation after the geographical range expansions induced by several catastrophic events, such as the Cretaceous mass extinctions and the glacial events of the Plio-Pleistocene (Veron 1995). Our finding of incongruity in the phylogenetic patterns among mitochondrial genes versus nuclear genes in *Stylophora*, are therefore consistent with multiples studies in reef building corals.

In addition, to our knowledge, the degree of incongruence in mtDNA genes found in our analyses has not being identified in others coral genera so far. In *Stylophora*, incongruence between the mt*ORF* and the *CR* topologies was reported by Arrigoni et al. (2016) and Stefani et al. (2011). These authors suggested that the most plausible hypothesis for explaining this finding was the presence of pseudogenes at the mitogenome of this group, but we found no evidence for stop codons or other signs of ORF disruption, rather evidence that this mt*ORF* may code for a functional transmembrane protein (Banguera-Hinestroza et al. unpublished results) as proposed by Flot and Tillier (2007). Therefore, we believe that other hypotheses such as recombination resulting from introgressive hybridization at the mtDNA genome of *Stylophora* are equally probable as the ones discussed in Stefani et al. (2011).

In species with a uniparental inheritance of mtDNA, genes are linked and are expected to show the same phylogenetic patterns (Ladoukakis and Zouros 2017). However, contrasting topologies in mtDNA genes may arise as the result of mtDNA introgressive hybridization, which is known to have a clear impact in genealogical and phylogenetic reconstructions (White et al. 2008). The presence of divergent mitochondrial genomes in the germline of an organism (heteroplasmy) as the result of paternal leakage (Rokas et al. 2003; Barr et al. 2005; Kuijper et al. 2015) may result in mtDNA recombination (considering that heteroplasmy have remained without modifications in the oocyte cytoplasm long enough for allowing recombination between paternal and maternal mtDNA; White et al., 2008). In the case that the recombinant genome become fixed, viable, and passed to the following generations it would result in viable hybrids carrying a mixed mtDNA genome (Rokas et al. 2003; White et al. 2008; Wilton et al. 2018). Even though, heteroplasmy have not been recorded in reef-building corals, there is an increasing evidence that mtDNA recombination resulting from mtDNA introgressive hybridization is not rare in nature (Rokas et al. 2003; Piganeau et al. 2004; Barr et al. 2005; Levsen et al. 2016).

### 3.2. mtDNA recombination explains topological incongruence

Our recombination analyses hypothesized the *RS_LinA* as a recombinant sequence with high probabilities in all methods (*P*-values ranging between 4.0 × 10^−03^ and 2.96 × 10^−29^) with the major parent being the sequence from *RS_LinB*, and only the minor parent the sequences from *S. pistillata* from Taiwan. For *RS_LinA* the recombinant region –inherited from the minor parent – spans the full *nd6*, *atp6* and mt*ORF* (Supplementary Figure *S3*), which explain that trees constructed with these genes showed a common ancestry between the *RS_LinA* and the Indo-Pacific/Indian ocean/Madagascar lineages, while the same trees show a strong diversification between these two lineages and the *RS_LinB* (Figure 2 and Figure 3). Moreover, the *RS_LinA*, inherited most of its mtDNA genes from the ancestor of *RS_LinB*. Therefore, it is expected that other mtDNA genes display a closest relationship between the *RS_LinA* and the *RS_LinB*, as shown in our phylogenies for the CR and 12S genes and in the mitogenome phylogenies excluding the recombinant region (Figure3b). Consequently, our data strongly supports mtDNA introgressive hybridization followed by recombination as the main cause of mito-mito topological incongruence in *Stylophora*.

We interpret these results as a strong evidence that *RS_LinA* likely arose from a hybridization event between two ancestral *Stylophora* lineages. The introgression pattern follow that recorded in species of hybrid zones, in which hybrids usually display an unequal mix of two mtDNA genetic backgrounds, with most mitochondrial genes or genetic polymorphism passed from the genetic pool of the local species to the genome of the “invader” species (asymmetric patterns of introgression; Harrison and Larson, 2014).

### 3.3. Interspecific phylogenies and comparative phylogeographic analyses of Red Sea lineages

As our study focuses at the population level –based on the three most variable genes used here: the *hsp70*, the CR and the *mtORF–* markedly different phylogeographic patterns arises for *RS_LinA* and for *RS_LinB* along the latitudinal and environmental gradient in the Red Sea (Figure 5). These patterns show a strong structure for *RS_LinB*, in which the groups supported by the phylogenetic analyses (Figure 4) have restricted distribution either in the northern Red Sea or in the southern Red Sea, with the break between the two zones found around Yanbu, a trend that was supported by the distribution of the haplotype frequencies along the gradient for each gene (Supplementary Figure *S2*). In contrast, a lack of association with northern and southern environmental provinces was found in members of *RS_LinA*, and rather a clinal variation in haplotype frequencies was found across the gradient (Supplementary Figure *S2*).

Both lineages were also remarkably different in terms of morphology and *Symbiodinium* association: in the northern Red Sea population of *RS_LinB* (i.e. subclade *LB1*; Figure 4b) most colonies were associated with morphotypes of *Stylophora subseriata* (as described in Veron et al. 2018) and *Symbiodinium* type C160; whereas the central-southern population (i.e. subclade LB2; Figure 4b) grouped mostly individuals described in the literature as *Stylophora pistillata* and were associated mainly with *Symbiodinium* type A1, but in contrast to the northern group, no colonies were found carrying C160 or A1-C160 *Symbiodinium* types. Contrary to the trends detected in *RS_LinB*, colonies belonging to the hybrid lineage (*RS_LinA*) showed high phenotypic plasticity and most colonies carried a *Symbiodinium* type A1 in both northern and southern regions (Figure 4).

The strong differences found in both lineages, in terms of phylogeography, population subdivision, and congruence/incongruence between morphology and phylogenies, reinforces the hypothesis of an early colonization of the Red Sea by members of *RS_LinB* and a recent colonization of *RS_LinA* in agreement with its hybrid origin. This evolutionary scenario could be explained on the light of the multiple processes that have affected the region during the last glacial periods, as well as of the demographic history of reef coral fauna in this region. Several studies targeting the geological dynamic of the Red Sea and the patterns of diversity and endemism of its fauna (Moustafa and Hallock 2008; Rohling et al. 2008; Fine et al. 2013; Moustafa et al. 2014; DiBattista, Howard Choat, et al. 2016; DiBattista, Roberts, et al. 2016) indicate that early coral reefs inhabiting this region (i.e. likely present in the region since the Miocene; Taviani 1998) were able to survive extreme climatic variations in sea levels and periods of hyper salinity (Rohling et al., 2013; Siddall et al., 2003; Righton et al., 1996; Rohling et al., 2013) thanks to the presence of refuge areas in the northern Red Sea (i.e. the Gulf of Aqaba) and in the southern regions (i.e. Southern Red Sea and Gulf of Aden) (Fine et al. 2013; Moustafa et al. 2014; Pellissier et al. 2014). Therefore, isolation in these refuge zones leading to diversification and posterior recolonization may explain the differentiation in allopatry of northern and central-southern populations of *RS_LinB*.

Furthermore, the phylogeographic patterns, such as the one presented here are also in accordance with several biogeographic studies, showing that oceanographic conditions along the environmental gradient of the Red Sea (Raitsos et al. 2013; Sawall et al. 2015) have originated barriers to genetic flow, influenced strongly the population structure of several species, and shaped patterns of endemism in this region (Nanninga et al. 2014; Saenz-Agudelo et al. 2015; DiBattista, Howard Choat, et al. 2016), despite the fact that they seem to have little impact on reef communities in terms of richness of species and species diversity (Roberts et al. 2016). For example, Nanninga et al. (2013) found a good correlation between the genetic differentiation among anemonefish (*Amphiprion bicinctus*) along the Saudi Arabian coast and the environmental heterogeneity in this basin, with the steadiest genetic break found at approximately 19°N (Southern Red Sea), which coincided with a sharp increase in turbidity in this zone. Results of our study also concur with those of Froukh and Kochzius (2008, 2007) who also found high genetic differentiation that were related to physical differences among fourline wrasse fish (*Larabicus quadrilineatus*) from the northern and southern Red Sea. On a larger scale, research on butterfly and angel fish by Roberts et al. (1992) and studies by DiBattista et al. (2013) have hypothesized that several reef fish species diversified and remained restricted to the Red Sea.

### 3.4. The diversification of *Stylophora* in the Red Sea and Gulf of Aden

Our phylogeographic analyses indicate a broad distribution of *RS_LinB* across the Red Sea but limited distribution in the Gulf of Aden (only few 5 out of 37 samples carrying a single mtDNA haplotypes were found in this region, see also Stephani et al. 2017) (Supplementary Figure *S2*), which suggest that the ancestor of *RS_LinB* was likely endemic of these regions, with the border of its distribution in the Gulf of Aden. This lineage likely interbred with a lineage of broader distribution in the Indo-Pacific/Indian-Ocean at the periphery of its range, which is also a confluence between several marine provinces: the Red Sea and Gulf of Aden, the western Indian Ocean, and Indo-Polynesian provinces, at zone that have been shown to hold several hybrid species (DiBattista et al. 2015). Moreover, the phylogenetic position of Arabian Gulf haplotypes, which were placed in the same subclades of the *RS_linA* specimens, implies that the hybrid expanded its range into the Arabian Gulf and into the Red Sea –where it lives in sympatry with the descendants of the parental lineage–

Our results concur with multiple studies that have identified a large number of hybrid species in the Red Sea (DiBattista, Howard Choat, et al. 2016; Berumen et al. 2017) and seem to fit in the model of hybrid zones and marginal areas (Barton 1979; Hewitt 1988), in which species from a diverse range of taxonomic groups (i.e. plants, yeasts and to a lesser extent animals) living at the edge of their geographic ranges produce viable hybrids carrying recombinant mitogenomes (Rokas et al. 2003; Barr et al. 2005). These hybrids are able to adapt to new climatic conditions, and diverge in sympatric association with their parental species (Rokas et al. 2003; Ballard and Whitlock 2004; Barr et al. 2005; Ujvari et al. 2007; Galtier 2011; Mastrantonio et al. 2016; Leducq et al. 2017; Peris et al. 2017; Fourie et al. 2018).

In addition, the pattern found in our results in *Stylophora* were similar to those found in hybrid species of marine reef-fishes, found at the intersection of three main biogeographical regions: the Red Sea and Gulf of Aden, the western Indian Ocean and the Indo-Polynesian provinces (DiBattista et al. 2015). These authors identify 14 hybrid species based in molecular markers and morphological characteristics, the gene flow was unidirectional, and the parental species were found at the periphery of their distribution range, where they showed the lowest abundance. Interestingly, as in our work with *Stylophora*, the reported hybrids were found to be the result of intercrosses between endemic species, with restricted distribution range in the Red Sea and Arabian Gulf, with species broadly distributed in the Indo-Pacific, and the major parental species was at the limit of its distribution and almost absent from the Gulf of Aden area.

### 3.5. Implications for conservation in a climate change scenario

Under the current challenges that climatic change poses for coral reef ecosystems, particularly with the rising of oceanic temperatures above the limit threshold of most species (1°C above mean summer maximum) (Pandolfi et al. 2011; Hughes et al. 2018) evolutionary studies with a clear assessment of the demographic history of species and populations are of great relevance, particularly to understand the main mechanisms involved in adaptation and speciation in variable and extreme environments. Our study, coupling recombination analyses, phylogenetic and phylogeographic patterns, suggest that mtDNA genes may play a key role in adaptation to extreme environments in coral species.

Two genes, the *atp6* gene and the mt*ORF* gene showed high mutation rates and extreme variability in the *RS_LinB* and amino acid and structural changes were recorded mainly at the mt*ORF* protein (Banguera-Hinestroza et al. unpublished data). Two nucleotide changes at positions 104 and 187 in the *atp6* gene revealed multiple paths, resulting in replacement of polar amino acids by non-polar amino acids or vice versa (see results), which we speculate may have led to changes in protein characteristics and function as reported by other authors (see Betts and Russell 2007). Interestingly, the *atp6* protein is not only essential in the oxidative phosphorylation system, responsible for providing the energy required for multiple cellular functions (Saccone et al. 2006) but also is part of a group of genes that are essential in releasing and controlling the proliferation of deleterious oxygen radicals (ROS) in the mitochondria (Kühlbrandt., 2015; Wirth et al., 2016), which are known for causing oxidative damage to symbionts and corals during exposition to high temperatures, with the subsequent bleaching and dead of corals (Downs et al. 2002; Smith et al. 2005). This finding, therefore, support our hypothesis that the multiple mutational changes at the mt*ORF* and *atp6* genes played a key role in the adaptation of *RS_LinB* to the changing environment and habitat conditions of the Red Sea.

In addition, in *RS_LinB*, the frequencies of the most common haplotypes of the mt*ORF* locus were associated with coldest (north) or hottest (central-south) areas in the Red Sea (Supplementary Figure S2), a tendency that was also followed by the haplotype frequencies of the *hsp70* gene –a chaperone protein involved in heat stress response (Mayer and Bukau 2005; Kvitt et al. 2016)–, suggesting that both proteins may have played a similar role in the adaptation to temperature in *RS_LinB* variants. This premise was supported by the computational characterization of the mt*ORF* protein (Banguera-Hinestroza et al. unpublished) in which searches against approx. 95 million protein domains classified in the CATH data base –using the pDomTHREADER approach (Lobley et al. 2009)–, revealed that domains in the mt*ORF* protein of *RS_LinB*, have the highest matches with annotated domains involved in the structural integrity of a complex or its assembly within or outside a cell (CATH domain code: 1s58A00) and domains that play a role in cell-to-cell, cell-to-matrix interactions, and response to stress (1ux6A01) with high levels of confidence (0.0001 < *P* < 0.001).

Our findings suggest that selective pressures imposed by the extreme environmental variations in the Red Sea along multiple geological periods, may have had an impact in the mitogenome of this lineage, with consequences in adaptation and speciation, as have been found in other taxa evolving in a broad range of environmental conditions (Hill 2016; Hill 2017; Lamb et al. 2018). Even though the *RS_LinB* hybridized with an Indo-Pacific/Indian Ocean lineage, the resulting hybrid (*RS_LinA*) did not inherited the variation at the *nd6*, *atp6* and mt*ORF* genes, this open the question whether other mitochondrial genes or mtDNA mechanisms may play a role in hybrid speciation and adaptation, or whether the hybrid may be more vulnerable to strong temperature variation (i.e. nowadays the *RS_LinA* is very rare in the Arabian Gulf Area; Banguera-Hinestroza et al. unpublished results).

Taken together, our data indicate that hybridization in corals and the adaptation of coral hybrids species could be more complex than commonly expected, involving little known mechanisms implicated in mitochondrial metabolism and mitochondrial genome evolution, as have been previoulsy emphasized by Dixon et al., (2015) that demonstrated a strong maternal effect in heat stress tolerance. Assisted hybridization has been proposed as an alternative for conservation of coral reef via genetic rescue (Chan et al. 2018); however, little is known about the molecular mechanisms enhancing hybrid adaptation and speciation under extreme climatic conditions. A clear understanding of these mechanisms coupled with the demographic history of coral populations is therefore needed to pursue effective conservation plans.

## 4. MATERIAL AND METHODS

### 4.1. Coral Sampling

We sampled a total of 725 colonies of *Stylophora* at 25 locations along the Red Sea coast in 2011 and 2012 from the Gulf of Aqaba in the North (Maqna, 28° 31’ 34.20 N, 34° 48’ 14.30″ E) to Farasan Island in the South (16° 31’ 38.5″ N, 42° 01’ 54.09″ E). Our sampling design covered the entire environmental gradient (Temperature: ~21 in the north to ~33°C in the south; Salinity: 41 PSU in the north to 37 PSU in the south) of the Red Sea and the four oceanographic provinces as defined in Raitsos et al. (2013) (Figure 1 and Supplementary Table *S2*). Colonies were determined to belong to morphotypes of *S. pistillata*, *S. subseriata*, *S. danae*, *S*. *kuelhmanni* and *S. wellsi* (Figure 1). Identification was carried out with the assistance of Dr. J.E.N Veron (personal communication) and by comparing his photographic records of Red Sea samples with our photographs taken during coral sampling. Small coral nubbins (~1cm) were collected in 3-7 m depth from colonies 5 m apart from each other to minimize sampling of clones and were preserved at room temperature in salt-saturated DMSO buffer (Seutin et al. 1991; Gaither et al. 2011)

### 4.2. DNA extraction, PCR and sequencing

Coral nubbins were placed in approximately 400 µL of lysis buffer (DNeasy Plant Minikit; Qiagen, Hilden, Germany®) with sterile glass beads and beaten for 30 s at full speed in a Qiagen Tissue Lyser II (Qiagen, Hilden, Germany®). After cell lysis, DNA was extracted following the protocol recommended by the manufacturer. Five mitochondrial genes and four nuclear genes were amplified. First we barcoded our samples using the mt*ORF*, which is considered the most variable gene for Pocilloporidae (Flot and Tillier 2007) and a short fragment of the adjacent ATPase subunit 6 gene (*atp6*). After this initial barcoding step, the mitochondrial control region (CR), the cytochrome *c* oxidase subunit I (*cox1*) and the mitochondrial 12S and 16S ribosomal RNA (rRNA) genes were amplified for a subset of samples. A subset was also amplified for several nuclear genes: the heat shock protein (*hsp70*) gene, the internal transcribed spacers (ITS1 and ITS2) and the Plasma Membrane Calcium ATPase (*PMCA*) gene. The 16S, 12S and *cox1* were amplified with primers designed in this study. The primers we included and their respective references are listed in the Supplementary Table *S3*.

After amplification, samples were cleaned using the Exostar 1-step protocol (Ilustra®) and sequenced in both forward and reverse directions; 5-10 specimens of each haplotype were sequenced twice to confirm the accuracy of the sequences. The forward and reverse chromatograms were assembled and edited using Chromas-Pro software version 2.1.5.1 (Technelysium, Pty Ltd). Sequences were first aligned using MUSCLE (Edgar 2004) as implemented in MEGA version 7.0.26 (Kumar et al. 2016) and when the resulting alignment contained multiple gaps we used MAFFT’s FFT-NS-i (Slow; iterative refinement method) to improve the alignment (Katoh and Standley 2013). All alignments were confirmed and corrected by eye and sequences were trimmed to the same length for further analyses.

For 12S and 16S, we removed a small region in which a long stretch of homopolymers did not allow high confidence in the alignment. Nuclear genes were phased using the programs PHASE Version 2.1 (Stephens and Donnelly 2003) using the input files obtained in SeqPHASE (Flot 2010).

In the few cases when ITS1, ITS2, or *PMCA* chromatograms showed multiple double peaks (as expected when sequencing a mixture of DNA sequences of unequal length; Flot et al., 2006) the indels from these chromatograms were not considered for downstream analyses and only clean regions with double peaks were included in the phasing process. In addition to our samples from the Red Sea, we amplified and sequenced the 12S and 16S markers of selected specimens from Flot et al. (2011) from several geographic regions (New Caledonia, Japan, Philippines and Madagascar). Moreover, previously published mt*ORF* and CR sequences (Flot et al. 2011; Stefani et al. 2011) as well as *hsp70* sequences (Klueter and Andreakis 2013) from other geographic regions (i.e. Gulf of Aden and Indo-Pacific) as well as from the Red Sea (i.e. mt*ORF* sequences belonging to *S. mamillata*: Arrigoni et al., 2016) were downloaded from the NCBI database and included in our analyses (accession numbers for these samples can be found in the respective references). Finally, we also included *cox1* sequences from Keshavmurthy et al. (2013) that were downloaded from the Dryad Digital Repository: (doi:10.5061/dryad.n2fb2).

### 4.3. Identification of zooxanthellae (*Symbiodinium*)

*Symbiodinium* types were identified by the amplification of the ITS2 rDNA in a set of *Stylophora* samples from the Red Sea collected throughout the same gradient as described above (N=281). We used the DGGE-ITS2 protocol described in LaJeunesse (2002) with the primers and protocols recommended in LaJeunesse et al. (2003). DGGE gels, electrophoresis and PCR conditions are fully described in Arif et al. (2014). Sequences were processed using Chromas-Pro software version 1.7.5 (Technelysium, Pty Ltd) and phylogenetic assignments were built by comparisons with previously published ITS2 sequences in the GeoSymbio data base (Franklin et al. 2012) and the Todd LaJeunesse’s database (https://131.204.120.103/srsantos/symbiodinium/sd2_ged/database/views.php).

### 4.4. Sequence variation, phylogenetic and phylogeographic analyses

Number of haplotypes (h), polymorphic sites per gene, and shared haplotypes within and among Red Sea regions (Maqna, Al-Wajh, Yanbu, Kaust, Jeddah, Doga, Farasan and Gulf of Aden; Figure 1) as well as for other geographic locations (see above), were calculated using Arlequin version 3.5 (Excoffier and Lischer 2010) and DNAsp version 5.1 (Librado and Rozas 2009). Phylogenetic analyses were performed using *Stylophora* specimens from seven geographic regions (Caledonia, Japan, Philippines, Madagascar, Gulf of Aden, Arabian Gulf and Red Sea). When several sequences shared the same haplotype, a single representative was included in the data set for phylogenetic analyses.

To account for the phylogenetic signal in indels (insertion-deletion polymorphism) gaps were treated as missing data and coded as additional presence/absence (0-1) characters with the program Fastgap v1.2. (Borchsenius 2009) resulting in a data set with two data partitions: nucleotides and standard characters (0-1). This approach was used to infer Bayesian trees with a mixed model, which allows the combination of different data partitions with parameters unlinked across partitions (Huelsenbeck and Ronquist 2001). The best evolutionary models for each gene and data partition were selected by calculating their BIC (Bayesian Information Criterion) scores as recommended in MEGA v.7 (Kumar et al. 2016). The non-uniformity of evolutionary rates among sites was modelled using a discrete Gamma distribution (+G) with 5 rate categories and by assuming a portion of the sites to be invariant (+I). The gamma shape parameter, proportion of invariant sites, transition/transversion ratio, nucleotide frequencies and rates of substitutions were also estimated from the data.

Bayesian trees were built in MrBayes (Huelsenbeck and Ronquist 2001) at the CIPRES Science Gateway v 3.1 (Miller et al. 2010). For each data set we performed 4 independent runs (nruns=4) with 1 cold and 3 heated chains (nchains=4). The total number of generations (ngen) was set at 10,000,000; sampling was performed every 1000 generations (Diagnfreq=1000); and the burn-in fraction was set to 25% (burninfrac=0.25). Convergence was assessed using the program Tracer v 1.7 (Rambaut and Drummond 2009). Maximum Likelihood (ML) and Neighbor Joining (NJ trees) were built in MEGA v.7 (Kumar et al. 2016), with 1000 bootstrap replicates. We used the nearest neighbor interchange algorithm (NNI) with an initial tree automatically generated (option: NJ/BioNJ) and the branch-swap filter set to strong, including all sites in the alignment.

NJ trees were constructed using the Maximum Composite Likelihood method, with the pairwise deletion option activated. Trees were constructed for all markers, except 16S and *cox1* that showed little or no variability in *Stylophora* spp. (see results). mt*ORF* and CR phylogenies (using our complete data set) were built only using the alignable regions among all samples and the outgroups (*Pocillopora* and *Seriatopora*), and alternatively, excluding the outgroups to allow the analyses of longer fragments (in these cases, the root of the trees was placed in agreement with that inferred by the phylogeny using shorter fragments and by the loci in which the full outgroup sequences were included).

Relationships among haplotypes were explored for the three most variable markers (mt*ORF*, CR and *hsp70*) using the program HaplowebMaker (https://eeg-ebe.github.io/HaplowebMaker/; Spöri and Flot, in prep.) applying the median-joining algorithm. Haplotypes belonging to specimens from the Pacific regions and Madagascar were included, when possible, to identify patterns of colonization. The frequency of each haplotype along the environmental gradient of the Red Sea (i.e. each locality/population) was calculated using Arlequin v. 3.5 (Excoffier and Lischer 2010).

### 4.5. Mitogenomes assembly and annotation

The complete mitogenomes of the two Red Sea lineages identify in our individual phylogenies analyses (see results) –called *RS_LinA* and *RS_LinB–* were obtained using two different approaches. First, paired-end Illumina reads from the draft genome of *Stylophora pistillata* (Voolstra et al. 2017) –identified as *RS_LinA–* were downloaded from the NCBI Short Read Archive (https://www.ncbi.nlm.nih.gov/sra/SRX999949) using fastq-dump (fastq-dump -- origfmt --split-3 SRR1980974). Second, 1.7 Gb paired-end reads for the *RS_LinB* were obtained from Red Sea samples preserved in CHAOS buffer (Flot 2007) and extracted using the DNA NucleoSpin kit (Macherey-Nagel) following the protocol recommended by the company. Sequencing was performed in an Illumina NovaSeq 6000 at the BRIGHTcore facilities (Brussels Interuniversity Genomics High Throughput core)

The mitogenome of both lineages were assembled de novo from the raw Illumina reads using NOVOPlasty v2.7.1 (Dierckxsens et al. 2016) with the *cox1* gene as a seed. For the assembly of the *RS_LinB* mitogenome, both *cox1* and the complete mitogenome of the *RS_LinA* were used as seeds. Runs were performed without and with reference, using the mitogenome of *Stylophora pistillata* (NC_011162.1) for comparative purposes. The accuracy of the assembly was confirmed by mapping the reads back to the assembled genomes using Bowtie2. In addition, the identity of each lineage was corroborated by aligning the resulting genomes with sequences from six of the markers used for our interspecific phylogenies (mt*ORF*, CR, *atp6*, 12S, 16S and *cox1*). Genes were annotated using MITOS (Bernt et al. 2013; Al Arab et al. 2017). Alignment for each annotated gene was performed with MUSCLE (Edgar 2004) and genes identities were confirmed using BLAST searches. When there was incongruence, particularly at the 5’ and 3’ ends of the genes, annotations were manually corrected using as reference standard (RefSeq) the mitogenome of *Stylophora pistillata* from the NCBI data base (NC_011162.1). Finally, genes were concatenated using Fastgap v1.2. (Borchsenius 2009) and aligned with mitogenomes from other members of the family Pocilloporidae using MAFFT (Katoh and Standley 2013). The mitogenomes included were: *Seriatopora hystrix* (EF633600.2), *Seriatopora caliendrum* (EF633601.2), *Pocillopora damicornis* (EF526302.1; EU400213.1), *Pocillopora eydouxi* (EF526303.1), *Stylophora pistillata* (of Indo-Pacific origin; NC_011162.1) and *Madracis mirabilis* (EU400212.1).

### 4.6. Analyses of recombination and mitogenome phylogenies

Recombination was tested using the alignments of the full mitogenomes from members of the family Pocilloporidae, including the two divergent *Stylophora* lineages from the present study – see results–; the genus *Polyciathus* was also included as an external sequence. Seven heuristic recombination detection methods implemented in the RDP4 package (Martin et al. 2015) were used: RDP (Martin et al. 2010), BOOTSCAN (Martin et al. 2005), GENECONV (Padidam et al. 1999), MAXCHI (Maximum Chi Square method; Smith 1992); CHIMAERA (Posada and Crandall 2001), SISCAN (Sister Scanning method; Gibbs et al. 2000), and 3SEQ (Boni et al. 2006).

The default settings were used for most methods, except for window sizes, step sizes and tree building methods. Window size was selected to include at least 10 variable nucleotide positions within every window examined. In RPD, SISCAN, and BOOTSCAN methods windows size were set at 100; in CHIMAERA and MAXCHI at 200. Alternatively, the MAXCHI and CHIMAERA methods were set to run with variable windows sizes, to allow the program to adjust the window size depending on degrees of parental sequence divergence as recommended by the author. Only recombination events detected with P<0.05, after Bonferroni correction were recorded.

The boundaries of the breakpoints were double checked by comparing the agreement of the boundaries between methods. The major parent and the minor parent hypothesized by the program, were contrasted against evidence from our phylogenetic and phylogeographic analyses. Furthermore, given that the parental ascription of the recombinant sequence may be difficult among closely related sequences, we refined the hypothesis by comparing the hypothesized distribution range of the major *Stylophora* clades (Keshavmurthy et al. 2013) and the supporting data from previous studies in hybrids species in the region and outside the Red Sea (DiBattista et al. 2015). It is important to note that the identified major and minor parents are not the “real” parental sequences, but sequences that are closely related to likely ancestral sequences, also that the recombinant region is related to the “minor” parent and the “non-recombinant” region related to the “major” parent (Martin et al. 2015).

Phylogenetic relationships among mitogenomes were inferred using maximum likelihood trees in PhylML (Guindon et al. 2010) available at the http://www.phylogeny.fr/index.cgi (Dereeper et al. 2008; Dereeper et al. 2010). Branch support was evaluated using an Approximate Likelihood Ratio test (aLRT) (Anisimova and Gascuel 2006). Trees were built for: (a) the full alignments, including and excluding the recombinant region and (b) genes within the recombinant region. The best model was selected as indicated in MEGA v7 (Kumar et al. 2016) as the GTR+G+I (General Time Reversible mode, non-uniformity of evolutionary rates among sites modeled using a discrete Gamma distribution with 5 rate categories and a fraction invariable sites).

Because ambiguously aligned regions or highly divergent sequences can mislead recombination signals, we ran the analyses including and excluding the *mtORF* and the *atp8* gene, and also excluding ambiguously aligned regions within the mt*ORF*. Finally, to discard the influence of highly divergent sequences, we repeated the recombination analyses excluding the mitogenomes of *Madracis* and *Polyciathus*. After analyses, polymorphic sites in hypervariable genes within the recombinant region for each *Stylophora* lineage were calculated using DNAsp version 5.1 (Librado and Rozas 2009).

## Acknowledgments

The authors want to thank Dr. Veron for assistance with voucher identification, Dr. Saenz Agudelo for sample collections and recommendations, Dr. Bouwmeester for sample collection, Vanessa Robitzch for helping with the DNA extractions, Catalina Ramirez for her support in bioinformatics pipelines and Sebastien Santini - CNRS/AMU IGS UMR7256, for its effort in maintaining the excellence of the Phylogeny.fr site. Thanks also to Sandra Cervantes Arango, Dario Ojeda Alayon for useful discussions and advices during the preparation of this manuscript. We thanks to King Abdulaziz University (KAU) and the Bioscience Core Lab at KAUST for sharing their facilities.

---

**Supplementary Table S1.** Sampling sites in the Red Sea and geographic coordinates

| *Locations* | *Reef* | *<sup>a</sup>Ns* | *<sup>b</sup>Nt* | *<sup>c</sup>Nseq* | *Coordinates* |  | *Depth* |
| --- | --- | --- | --- | --- | --- | --- | --- |
| MAQNA | R1 | 30 | 67 | 67 | 28° 31' 34.20" | 34° 48' 14.30" | 3-5 |
|  | R2 | 37 |  |  | 28° 25' 36.48" | 34° 45' 06.17" | 3-7m |
| AL-WAJH | R1 | 59 | 113 | 113 | 26° 11' 15.05" | 36° 20' 57.21" | 1-3m |
|  | R2 | 2 |  |  | 26° 09' 59.80" | 36° 23' 32.20" | 3m |
|  | R3 | 24 |  |  | 26° 14' 28.3" | 36° 26' 25.3" | 5-10m |
|  | R4 | 28 |  |  | 26° 11' 06.2" | 36° 22' 58.9" | 3-7m |
| YANBU | R1 | 28 | 118 | 118 | 23° 56' 50.70" | 38° 10' 31.80" | 6-10 |
|  | R2 | 16 |  |  | 23° 54' 58.92" | 38° 09' 00.63" | 6-10 |
|  | R3 | 28 |  |  | 23° 57' 19.20" | 38° 12' 16.00" | 3-5m |
|  | R4 | 29 |  |  | 23° 54' 41.4" | 38° 09' 08.4" | 3-7m |
|  | R5 | 17 |  |  | ----- | ----- | 2-10m |
| KAUST | R1 | 42 | 135 | 135 | 22° 19' 09" | 38° 51' 16" | 1-7m |
|  | R2 | 46 |  |  | 22° 04' 02" | 38° 46' 09" | 2-8m |
|  | R3 | 47 |  |  | 22° 30' 48" | 38° 55' 17" | 2-10m |
| JEDDAH | R1 | 30 | 70 | 64 | 21° 45' 11.40" | 38° 57' 45.09" | 1-3m |
|  | R2 | 20 |  |  | 21° 46' 06.3" | 38° 57' 47.07" | 2-5m |
|  | R3 | 20 |  |  | 21° 41' 23.26" | 39° 00' 49.05" | N/A |
| DOGA | R1 | 29 | 88 | 88 | 19° 38' 06.40" | 40° 34' 31.30" | 3-5m |
|  | R2 | 29 |  |  | 19° 36' 50.50" | 40° 38' 17.50" | 3-5m |
|  | R3 | 30 |  |  | 19° 39' 56.49" | 40° 37' 21.59" | 3-5m |
| FARASAN | R1 | 30 | 134 | 131 | 16° 34' 45.50" | 42° 08' 57.50" | 4-5m |
|  | R2 | 30 |  |  | 16° 34' 44.38" | 42° 14' 11.47" | 3-5m |
|  | R3 | 20 |  |  | 16° 31' 30.65" | 42° 01' 57.11" | 3-4m |
|  | R4 | 30 |  |  | 16° 31' 38.5" | 42° 01' 54.09" | 3-6m |
|  | R5 | 24 |  |  | ----- | ----- | 1-5m |
| Total |  |  | **725** | **716** |  |  |  |

<sup>a</sup> = Total number of samples per reef; <sup>b</sup> = Total number of samples per region; <sup>c</sup> = Total number of sequences obtained for the mtORF gene

**Supplementary Table S2.** Polymorphic sites per gene

|  | ALL REGIONS |  |  |  |  | RED SEA |  |  |  |  |
| --- | --- | --- | --- | --- | --- | --- | --- | --- | --- | --- |
| Locus | Nseq | size (bp) | Polymorphic sites | Indels | Parsimony informative sites | Nseq | size (bp) | Polymorphic sites | Indels | Parsimony informative sites |
| **nDNA genes** |  |  |  |  |  |  |  |  |  |  |
| *ITS1* | 75 | 416 | 69 | 94 | 54 | 44 | 416 | 33 | 31 | 16 |
| *ITS2* | 56 | 329 | 90 | 22 | 68 | 42 | 329 | 44 | 3 | 26 |
| *PMCA* | 86 | 956 | 107 | 65 | 100 | 50 | 956 | 26 | 11 | 18 |
| *hsp70* | 738 | 732 | 119 | 2 | 70 | 646 | 732 | 91 | 2 | 33 |
| **mtDNA genes** |  |  |  |  |  |  |  |  |  |  |
| *atp6-mtORF (all samples)* | 827 | 545 | 108 | 99 | 107 | 714 | 545 | 102 | 63 | 102 |
| *atp6-mtORF (Clade 1)* | 461 | 884 | 48 | 291 | 40 | 353 | 884 | 10 | 255 | 9 |
| *Atp6-mtORF (Clade 2)* | 366 | 943 | 22 | 207 | 21 | 361 | 943 | 22 | 207 | 21 |
| CR | 401 | 757 | 68 | 146 | 65 | 325 | 757 | 3 | 146 | 3 |
| 12s | 72 | 1381 | 33 | 0 | 33 | 50 | 1381 | 6 | 0 | 5 |
| 16s | 61 | 1393 | 7 | 0 | 7 | 43 | 1393 | 1 | 0 | 1 |
| *COI* | 147 | 848 | 19 | 0 | 12 | 110 | 848 | 0 | 0 | 0 |

**Supplementary Table S3.**
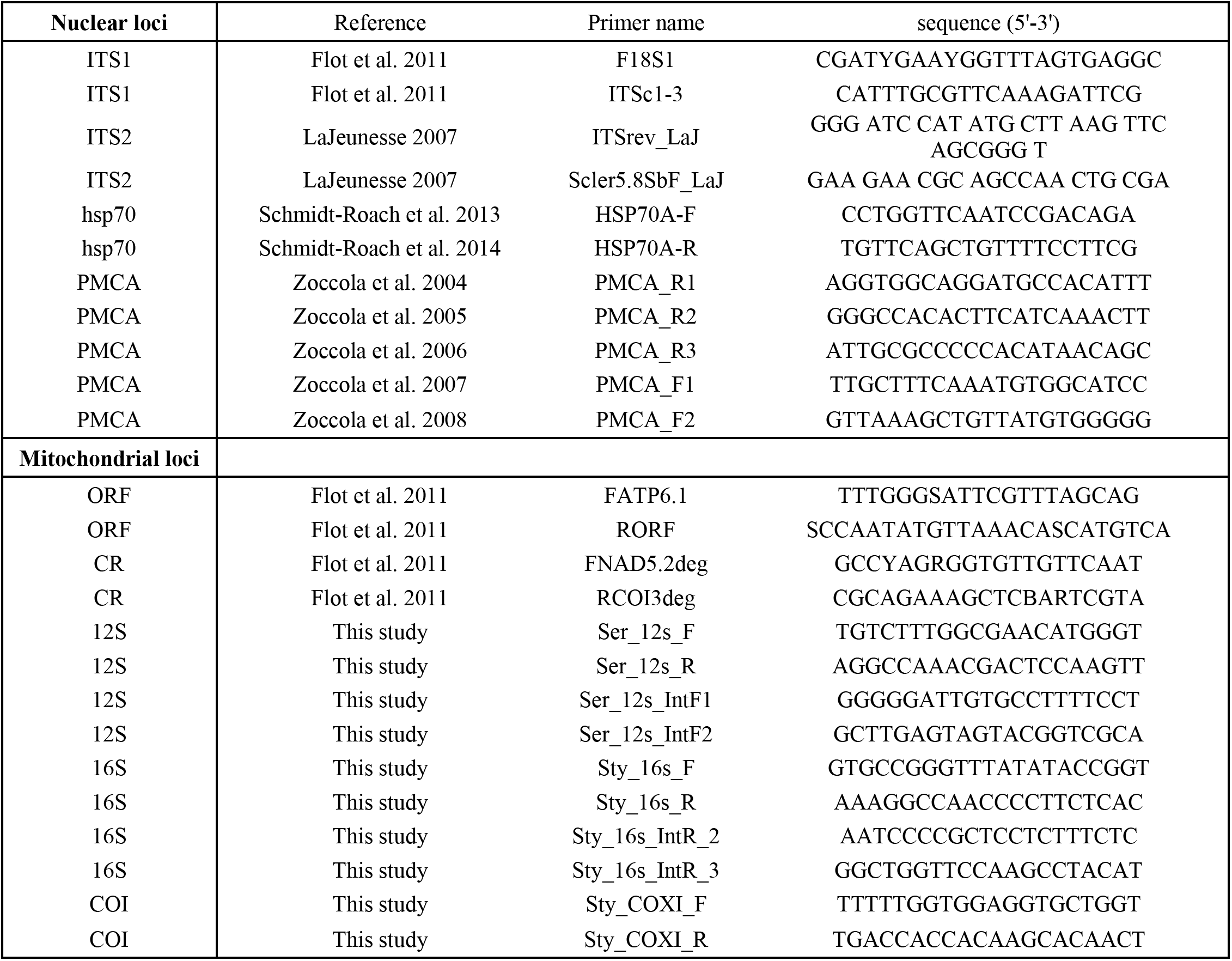
Primers and their respective references

**Supplementary Figure S1.**
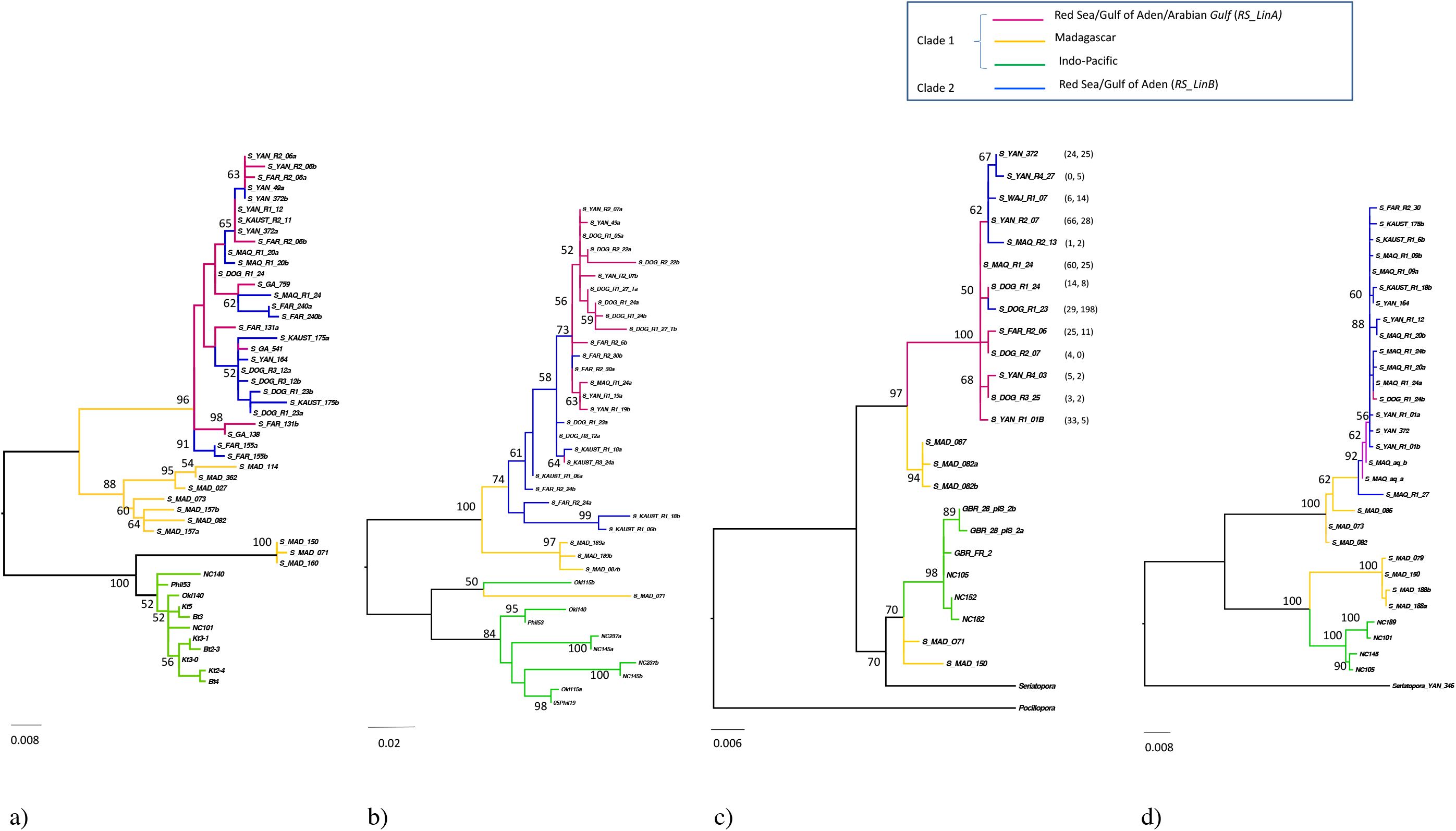
Nuclear phylogenies of the genus *Stylophora*. Bayesian trees for nuclear genes. **a)**. ITS1 b. **b)** ITS2 **c)**. *hsp70* **d)**. *PMCA.* Colors and haplotypes names are in agreement with those in the mt*ORF* phylogenies (see legend in Figure 2). In the *hsp70* phylogeny numbers in parenthesis correspond to the number of specimens found per haplotype in each lineage: *RS_LinA*, *RS_linB*.

**Supplementary Figure S2.**
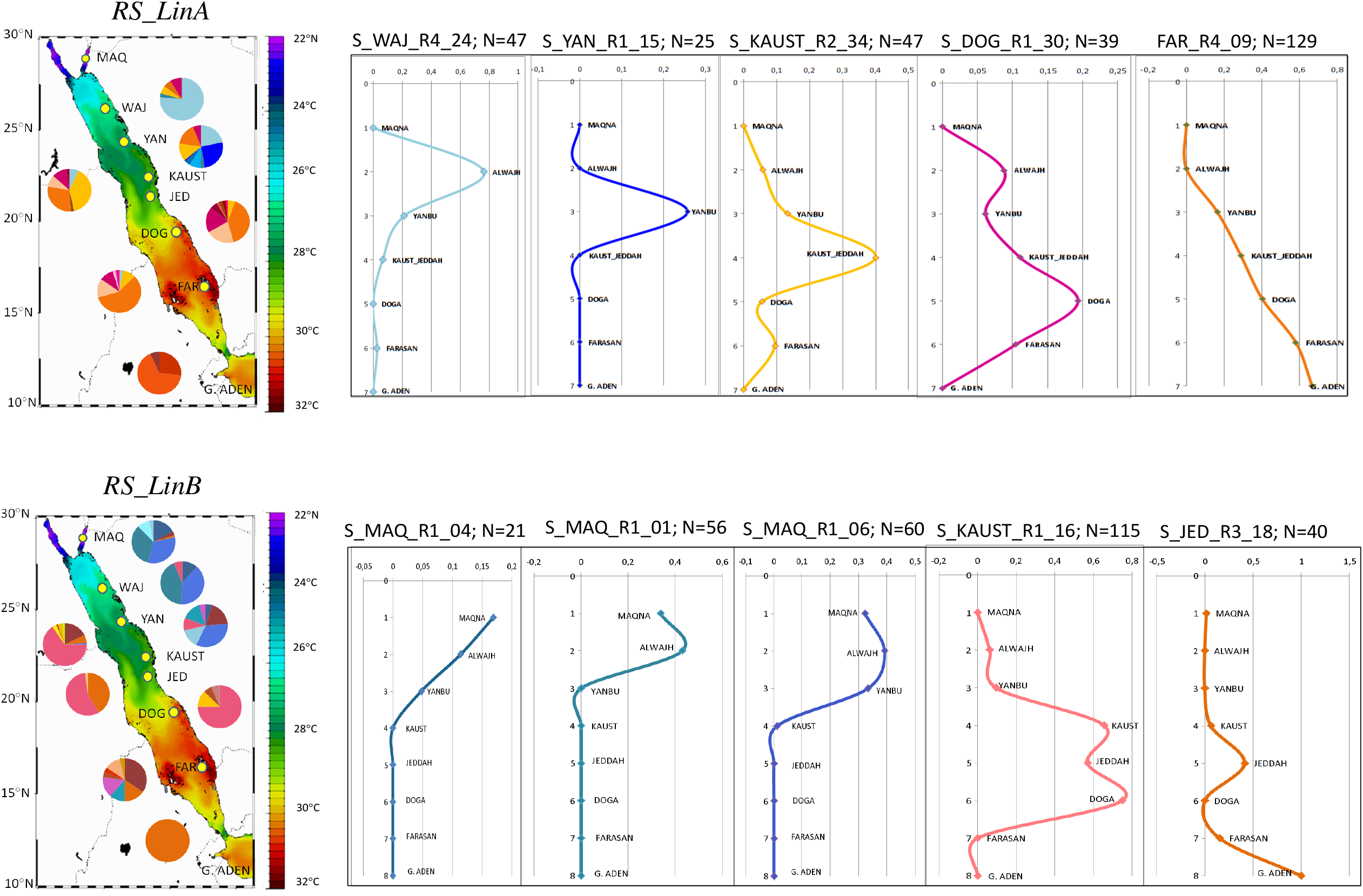
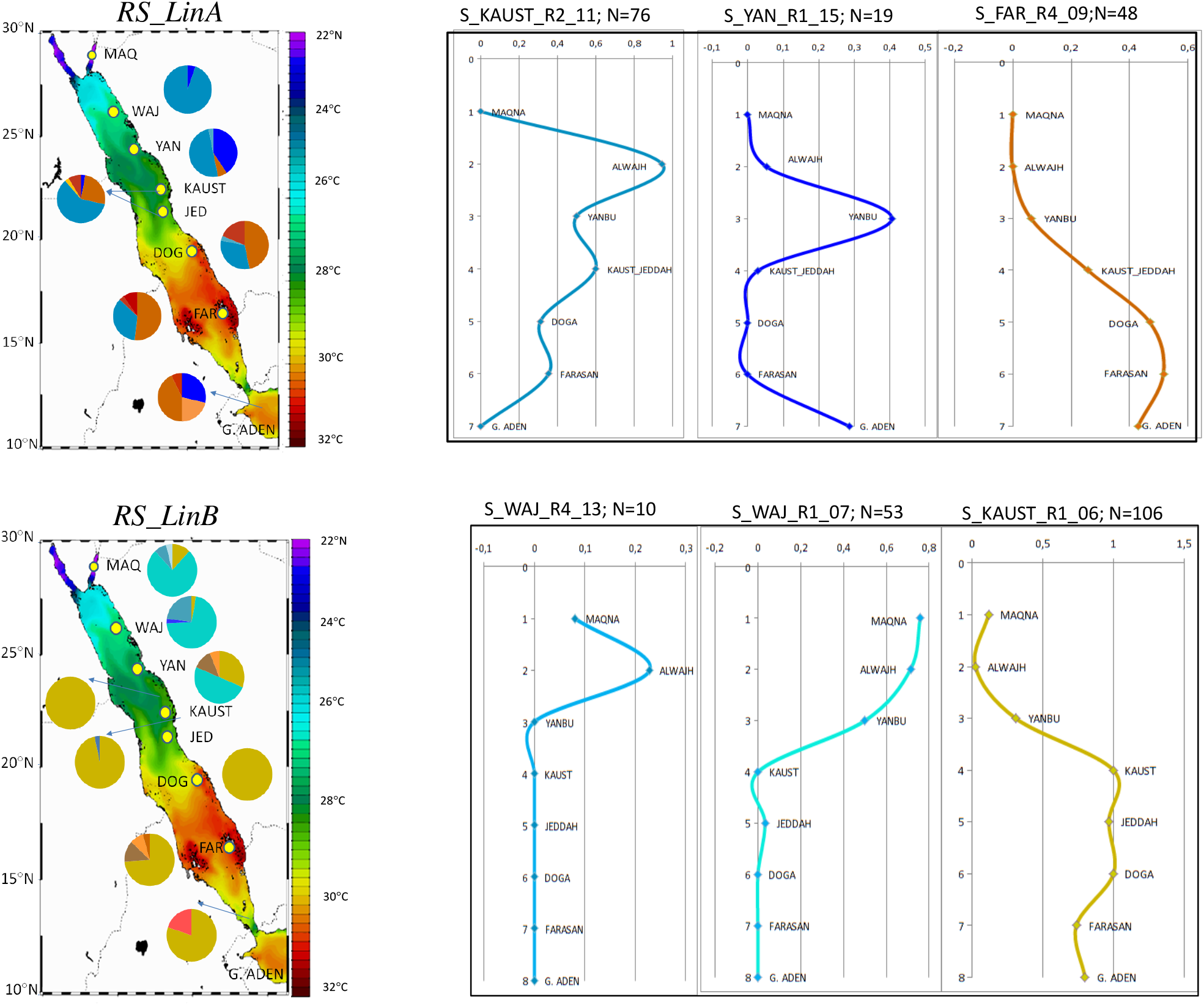
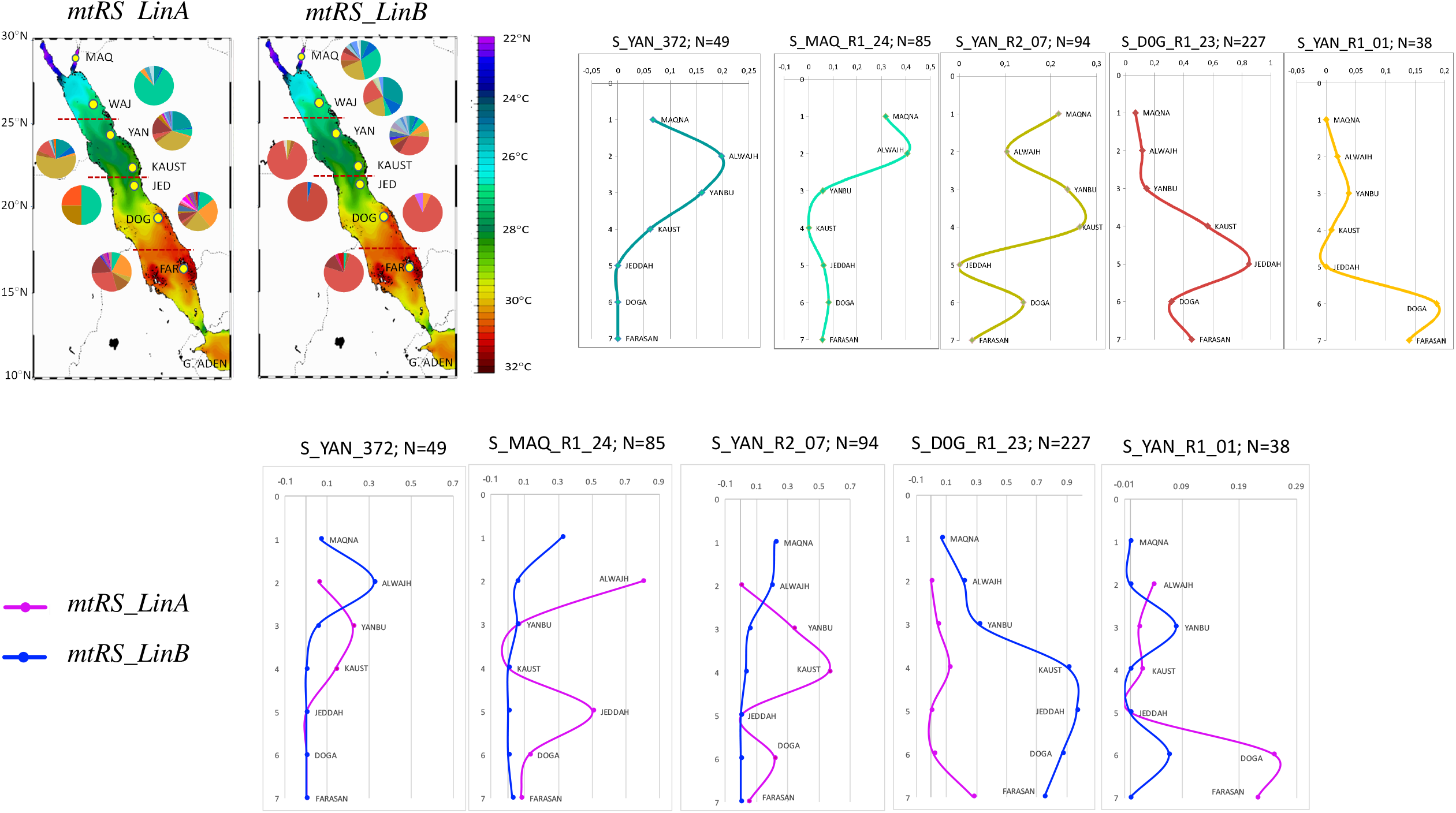
Distribution of haplotype frequencies for the most common haplotypes of the most variable genes across the environmental gradient of the Red Sea. Frequencies are indicated as per mtDNA lineage. The axis × represents relative frequencies. ***S2***a. ***atp6***-mt*ORF* locus ***S2***b. CR *gene*, and ***S2c***. ***hsp70*** gene, including all haplotypes (above) and dividing haplotypes per mtDNA lineage (below).

**Supplementary Figure S3.**
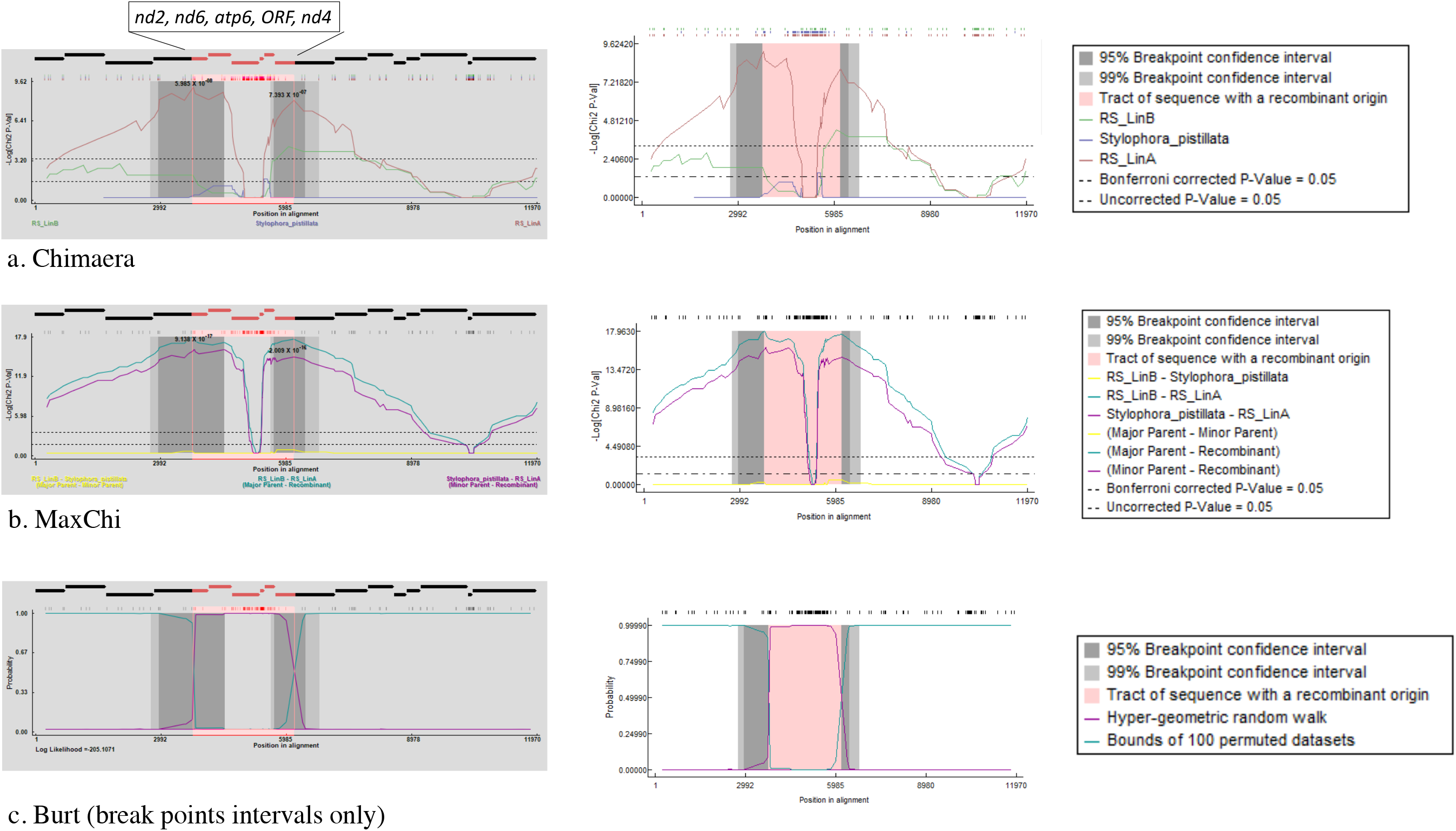
Identification of recombinant sequences and estimation of break points by different methods **a)**. CHIMAERA **b)**. MAXCHI, and **c)**. BURT

